# Role of Arginine and its Metabolism in TGF-β-Induced Activation of Lung Fibroblasts

**DOI:** 10.1101/2024.11.01.618293

**Authors:** Robert B Hamanaka, Kun Woo D Shin, M Volkan Atalay, Rengul Cetin-Atalay, Hardik Shah, Jennifer C Houpy Szafran, Parker S Woods, Angelo Y Meliton, Obada R Shamaa, Yufeng Tian, Takugo Cho, Gökhan M. Mutlu

## Abstract

Arginine is a conditionally essential amino acid with known roles in protein production, nitric oxide synthesis, biosynthesis of proline and polyamines, and regulation of intracellular signaling pathways. Arginine biosynthesis and catabolism have been linked to TGF-β-induced activation of fibroblasts in the context of pulmonary fibrosis; however, a thorough study on the metabolic and signaling roles of arginine in the process of fibroblast activation has not been conducted. Here, we used metabolic dropouts and labeling strategies to determine how activated fibroblasts utilize arginine. We found that arginine limitation leads to activation of GCN2 while inhibiting TGF-β-induced mTORC1 activation and collagen protein production. Extracellular citrulline could rescue the effect of arginine deprivation in an ASS1-dependent manner. Using metabolic tracers of arginine and its precursors, we found little evidence of arginine synthesis or catabolism in lung fibroblasts treated with TGF-β. Extracellular ornithine or glutamine were the primary sources of ornithine and polyamines, not arginine. Our findings suggest that the major role for arginine in lung fibroblasts is for charging of arginyl-tRNAs and for promotion of mTOR signaling.

**Highlights:**

- Arginine depletion inhibits TGF-β-induced transcription in human lung fibroblasts (HLFs).
- Arginine is not significantly catabolized in HLFs either through NOS or ARG dependent pathways.
- Extracellular glutamine and ornithine are the primary sources of polyamines in lung fibroblasts.
- The primary role of arginine in lung fibroblasts is for signaling through mTOR and GNC2.

## Introduction

Idiopathic Pulmonary Fibrosis (IPF) is a fatal disease, with a median survival of 3.5 years and affecting approximately 150,000 people in the United States [1; 2]. A defining feature of IPF is the Transforming Growth Factor-β (TGF-β)-dependent activation of lung fibroblasts, leading to the excessive secretion of extracellular matrix proteins, including collagen [3–6]. Lung fibroblasts are the primary cells responsible for the structural remodeling and impairment of lung function characteristic of IPF and thus represent a key therapeutic target for the treatment of the disease [6–8].

Metabolic reprogramming has emerged as a key regulator of fibroblast activation and is increasingly studied as a target of therapeutic intervention for IPF [9–12]. We have previously demonstrated that *de novo* synthesis of glycine and proline, the two most abundant amino acids present in collagen protein, is critical for collagen protein synthesis downstream of TGF-β [13–15]. How the metabolism of other amino acids is regulated during fibrotic processes is poorly understood.

Arginine is a conditionally essential amino acid that has been linked with fibrotic processes; however, the role of arginine in lung fibroblasts is poorly understood [16]. Arginine is catabolized by arginase enzymes (ARG1, ARG2), producing ornithine, which is a precursor for proline and polyamines. In the liver, where the urea cycle is active, ornithine can be converted to citrulline, which is then converted to argininosuccinate by argininosuccinate synthase (ASS1) followed by resynthesis of arginine by argininosuccinate lyase (ASL). While the complete urea cycle is not present in all cells, citrulline can also be produced directly from arginine through the activity of nitric oxide synthases (NOS) and be converted to arginine by ASS1 and ASL [17–21].

Both arginine synthesis and catabolism have been linked with lung fibrosis. ASS1 expression has been suggested to be reduced in IPF fibroblasts, causing these cells to be more dependent on extracellular arginine than control fibroblasts [22]. Arginine metabolites including ornithine, proline, and polyamines have been shown to be increased in the lungs of patients with IPF and in the fibrotic lungs of bleomycin-treated mice [23–25]. Arginase expression is increased in the lungs of bleomycin-treated mice and in lung fibroblasts and vascular smooth muscle cells (VSMCs) after treatment with TGF-β [26–28]. Chemical inhibition of arginase in lung fibroblasts has been shown to reduce collagen production after TGF-β, while inhibition of ornithine amino transferase (OAT), the enzyme which converts ornithine to proline inhibited TGF-β-induced collagen production in VSMCs and lung fibroblasts [26; 28; 29]. OAT expression correlates with %FVC decline in IPF patients and OAT inhibition was recently shown to inhibit development of fibrosis downstream of TGF-β [23; 29].

In addition to its central role in nitrogen metabolism, arginine is an important signaling regulator. We and others have demonstrated that the Mechanistic Target of Rapamycin Complex 1 (mTORC1) is a major regulator of amino acid homeostasis in lung fibroblasts [30; 31]. mTORC1 integrates signals from extracellular signaling cascades and intracellular nutrients, and is known to be particularly sensitive to cellular levels of arginine [32–35]. Arginine is also a negative regulator of the GCN2-mediated branch of the integrated stress response which responds to the accumulation of uncharged tRNAs, inhibiting protein translation [36].

Despite the continued interest on the role of arginine metabolism in fibrotic phenotypes, a thorough analysis of arginine metabolism in lung fibroblasts has not been conducted. Here, we used media drop out experiments and metabolic tracing to determine how TGF-β signaling regulates arginine biosynthesis and catabolism. We find that extracellular arginine is required for TGF-β-induced activation of mTORC1 and for collagen protein production. Arginine dependency was reduced when cells were cultured in Human Plasma Like Medium (HPLM) which contains the arginine precursors ornithine and citrulline. Addition of excess citrulline to the medium completely rescued the effect of arginine deprivation, suggesting that *de novo* biosynthesis can provide sufficient arginine in the absence of extracellular arginine. ASS1 was required for collagen production in the absence of extracellular arginine. While IPF fibroblasts contained lower cellular levels of arginine compared with control fibroblasts, we did not find a deficiency in arginine biosynthesis in these cells. We then performed metabolic labeling using arginine and its precursors. When cells were cultured in HPLM, we found little evidence of arginine synthesis or catabolism in lung fibroblasts. When cells were cultured in DMEM, we found increased arginine catabolism by both NOS and arginase; however, surprisingly, glutamine was the main source of cellular ornithine in these cells, contributing to proline and polyamine biosynthesis. Our findings suggest that arginine catabolism is dispensable for lung fibroblasts, and that the major role of arginine in lung fibroblasts is for charging of arginyl-tRNAs and for activation of mTORC1.

## MATERIALS AND METHODS

### Fibroblast Culture

Normal human lung fibroblasts and IPF lung fibroblasts (Lonza) were cultured in FGM2 (PromoCell) as previously described [13]. Cells were serum starved in DMEM (Gibco) containing 0.1% bovine serum albumin (BSA), 5.5mM glucose, 2 mM glutamine, and 1mM pyruvate for 24 hours prior to treatment with TGF-β (1ng/mL, Peprotech). For arginine starvation in DMEM, SILAC DMEM Flex Media (Gibco) was supplemented with 5.5mM glucose, 2mM glutamine, 1mM pyruvate, and 0.8mM lysine hydrochloride and 0.1% BSA. Arginine was added to control media at 0.4mM. For experiments with HPLM, media was formulated according to the original recipe protocol of Cantor et al [37] with the exception of arginine, citrulline, and ornithine. These were dissolved in Arg-Orn^−^Cit^−^ HPLM as 10x stocks and added to the parental media as indicated. Labeled metabolites were purchased from Cambridge Isotope Laboratories. ^13^C_6_ Arginine (CLM-2265), Guanido ^15^N_2_ Arginine (NLM-395), ^15^N_2_ Ornithine (NLM-3610), 4,4,5,5-D_4_ Citrulline (DLM-6039), ^13^C_5_ Glutamine (CLM-1822).

### siRNA Knockdowns

For siRNA knockdowns, 1×10^6^ NHLFs were transfected with 250 pmol ON-Silencer Select siRNA (Ambion). Cells were plated on 10cm dishes for 24 hours and then replated for experiments as above. Ambion product numbers are: nontargeting siRNA-4390843, siASS1 1-S1684, siASS1 2-S1685.

### Western Blotting

Cells were lysed, and electrophoresis was performed as we previously described [38]. Wells were lysed in 100μL Urea Sample Buffer (8M deionized urea, 1% SDS, 10% Glycerol, 60mM Tris pH 6.8, 0.1% pyronin-Y, 5% β-mercaptoethanol). Lysates were run through a 28 gauge needle and were electrophoresed on Criterion gels (Bio-Rad) and transferred to nitrocellulose using a Trans-Blot Turbo (Bio-Rad) set to the Mixed MW program. Primary antibodies used were: Collagen 1 (Abcam, ab138492), α-SMA (Sigma, A2547), Phospho-GCN2 (Cell Signaling, 94668), GCN2 (Cell Signaling, 3302), Phospho-P70S6K (Cell Signaling, 9234), P70S6K (Cell Signaling, 9202), Phospho-SMAD2/3 (Cell Signaling, 9520), SMAD2/3 (Cell Signaling, 9523), ASS1 (Invitrogen, PA-5-82740), ASL (Novus, NBP1-87462), NOS3 (Proteintech, 27120-1-AP), ARG2 (Invitrogen PA5-78820), GAPDH (Cell Signaling, 2118).

### RNA Isolation and Quantitative PCR

RNA as isolated using the GenElute Total RNA Purification Kit (Sigma) and reverse transcribed using iScript Reverse Transcription Supermix (Bio-Rad). Quantitative mRNA expression was determined by real-time RT-PCR using ITaq Universal SYBR Green Supermix (Bio-Rad). Primers used for PCR were: *COL1A1* (F:5’-GGTCAGATGGGCCCCCG-3’, R:5’-GCACCATCATTTCCACGAGC-3’), *ACTA2* (F:5’-GGCGGTGCTGTCTCTCTAT-3’, R:5’-CCAGATCCAGACGCATGATG-3’), *CTGF* (F:5’-GGCTTACCGACTGGAAGAC-3’, R:5’-AGGAGGCGTTGTCATTGG-3’), *SERPINE1* (F:5’-GGCTGACTTCACGAGTCTTTC-3’, R:5’-GCGGGCTGAGACTATGACA-3’).

### RNAseq Analysis

Total RNA isolated as above was sequenced on an Illumina NovaSEQ6000 at the University of Chicago Genomics Core Facility (100bp paired end). Sequence qualities of generated FASTQ files were assessed using FastQC. Transcript expression was quantified using Kallisto v.0.46.1 [39]. The Kallisto index was created with the GENCODE (Human Release 45, GRCh38.14) quantification was performed in quant mode using default parameters. Gene abundances were computed using the R package txiimport v.1.30.0 [40]. Differential gene expression analysis was performed using the edgeR package v.4.0.16 quasi-likelihood F model [41]. Differential gene expression was considered significant for genes with an FDR-adjusted p-value ≤0.05. Gene set enrichment analysis was performed on Hallmark pathways from MsigDB using the clusterProfiler package v.4.10.1 with R. All packages were run in RStudio (2023.06.2+561) with R version 4.3.1.

### Liquid Chromatography Mass Spectrometry

After treatment as above, cells were quenched with the 1 mL of dry ice cold 80% methanol and stored at −80°C until analysis. On the day of the analysis, samples were sonicated for 3 minutes in ice-cold water filled sonicator followed by 5 minutes of mixing with the thermomixer at 2000 rpm and 4°C, 20 minutes incubation on ice and centrifuge at 18,000g and 4°C for 20 minutes. The supernatant was dried down using the Genevac EZ-2.4 elite evaporator. The dried samples were resuspended in ice-cold 60/40 acetonitrile/water before LC-MS analysis. All solvents were LC-MS grade and obtained from Fisher Scientific.

Metabolites separation and detection was performed as described in [42]. In brief, Thermo Scientific Vanquish Horizon UHPLC system and Atlantis BEH Z-HILIC (2.1×150 mm, 2.5 µM; part # 186009990; Waters Corporation) column at acidic pH or iHILIC-(P) Classic (2.1×150 mm, 5 µm; part # 160.152.0520; HILICON AB) column at basic pH was used to detect the urea cycle related metabolites. For the acidic pH method, the mobile phase A (MPA) was 10 mM ammonium formate containing 0.2% formic acid and mobile phase B (MPB) was acetonitrile containing 0.1% formic acid. The column temperature, injection volume, and flow rate were 30°C, 5 µL, and 0.2 ml/minute, respectively. The chromatographic gradient was 0 minute: 90% B, 15 minutes: 20% B, 16 minutes: 20% B, 16.5 minutes: 90% B, 17 minutes: 90% B, and 23 minutes: 90% B. The flow rate was increased to 0.4ml/minute for 4.7 minutes during the re-equilibration. MS detection was done using Orbitrap IQ-X Tribrid mass spectrometer (Thermo Scientific) with a H-ESI probe operating in switch polarity mode for both methods except the in-vitro ^13^C_5_ citrulline tracing experiment data were collected only in positive mode. MS parameters were as follows: spray voltage: 3800 V for positive ionization and 2500 V for negative ionization modes, sheath gas: 80, auxiliary gas: 25, sweep gas: 1, ion transfer tube temperature: 300°C, vaporizer temperature: 300°C, automatic gain control (AGC) target: 25%, and a maximum injection time of 80 milliseconds (ms). For the basic pH method, MPA was 20 mM ammonium bicarbonate at pH 9.6, adjusted by ammonium hydroxide addition and MPB was acetonitrile. The column temperature, injection volume, and the flow rate were 40°C, 2 µL, and 0.2ml/minute, respectively. The chromatographic gradient was 0 minute: 85% B, 0.5 minute: 85% B, 18 minutes: 20% B, 20 minutes: 20% B, 20.5 minutes: 85% B and 28 minutes: 85% B. MS parameters were as follows: spray voltage:3600V for positive ionization and 2800 for negative ionization modes, sheath gas: 35, auxiliary gas: 5, sweep gas: 1, ion transfer tube temperature: 250°C, vaporizer temperature: 350°C, AGC target: 100%, and a maximum injection time of 118 ms.

For both methods, data acquisition was done using the Xcalibur software (Thermo Scientific) in full-scan mode with a range of 70-1000 m/z at 120K resolution (acidic pH) and 60K (basic pH). Metabolite identification was done by matching the retention time and MS/MS fragmentation to the reference standards. Data analysis was performed using Thermo Scientific Tracefinder 5.1, Compound Discoverer 3.3 software and natural abundance correction was performed using the IsoCor [43]

### Analysis of scRNAseq Data sets

Preanalyzed data from Habermann *et al* [44] were downloaded from GitHub (https://github.com/tgen/banovichlab/tree/master/pulmonary_fibrosis/10x_scRNA-Seq_2019). This dataset is also publicly available at GEO: GSE135893. scRNA-seq data were processed and analyzed using Seurat v5 package in R version 4.4.0. We excluded cells with fewer than 250 detected genes or larger than 20% mitochondrial genes. Seurat v5 was used to perform dimensionality reduction, clustering, and visualization. Recursive clustering analysis of subpopulations was conducted to improve the granularity of cell annotations and to obtain fibroblast cells. Cell-type annotation of fibroblasts was performed based on markers defined by Habermann. Visualization of the cells and clusters on a 2D map was performed with uniform manifold approximation and projection (UMAP). Dot plots and UMAP plots overlaid with gene expression levels were generated using Seurat.

### Statistical analysis

qRT-PCR and metabolomic data were analyzed in Prism 10 (GraphPad Software, Inc). All data are shown as mean ± standard error of the mean (SEM). Significance was determined by one-way or two-way ANOVA using Tukey’s correction for multiple comparisons. * P < 0.05, ** P < 0.01, *** P < 0.001.

## Results

### Arginine is required for TGF-β-induced signaling and collagen production

To determine how arginine regulates fibroblast signaling and activation downstream of TGF-β, we cultured cells in DMEM containing standard arginine concentration (0.4mM) or without arginine and treated with TGF-β for 0, 24, or 48 hours (**Fig. 1A**) We found that arginine deficiency inhibited TGF-β-induced induction of α-smooth muscle actin (α-SMA) and greatly reduced induction of collagen protein. This was associated with reduced phosphorylation of the mTOR target S6-kinase consistent with the role of arginine as an mTOR activator. We also observed phosphorylation of GCN2, suggesting that arginine deprivation leads to accumulation of uncharged arginyl-tRNAs in HLFs. To determine how arginine deficiency regulated canonical signaling downstream of TGF-β, we measured phosphorylation of SMAD2/3 in the presence or absence of arginine. We found that SMAD phosphorylation was attenuated by arginine deficiency (**Fig. 1B**). Consistent with this, when we examined expression of SMAD target genes, we found reduced TGF-β-induced expression of *COL1A1*, *ACTA2*, *CTGF*, and *SERPINE1* in arginine deficient cells suggesting that arginine is required for the transcriptional responses to TGF-β in lung fibroblasts (**Fig. 1C**).

**Figure 1.**
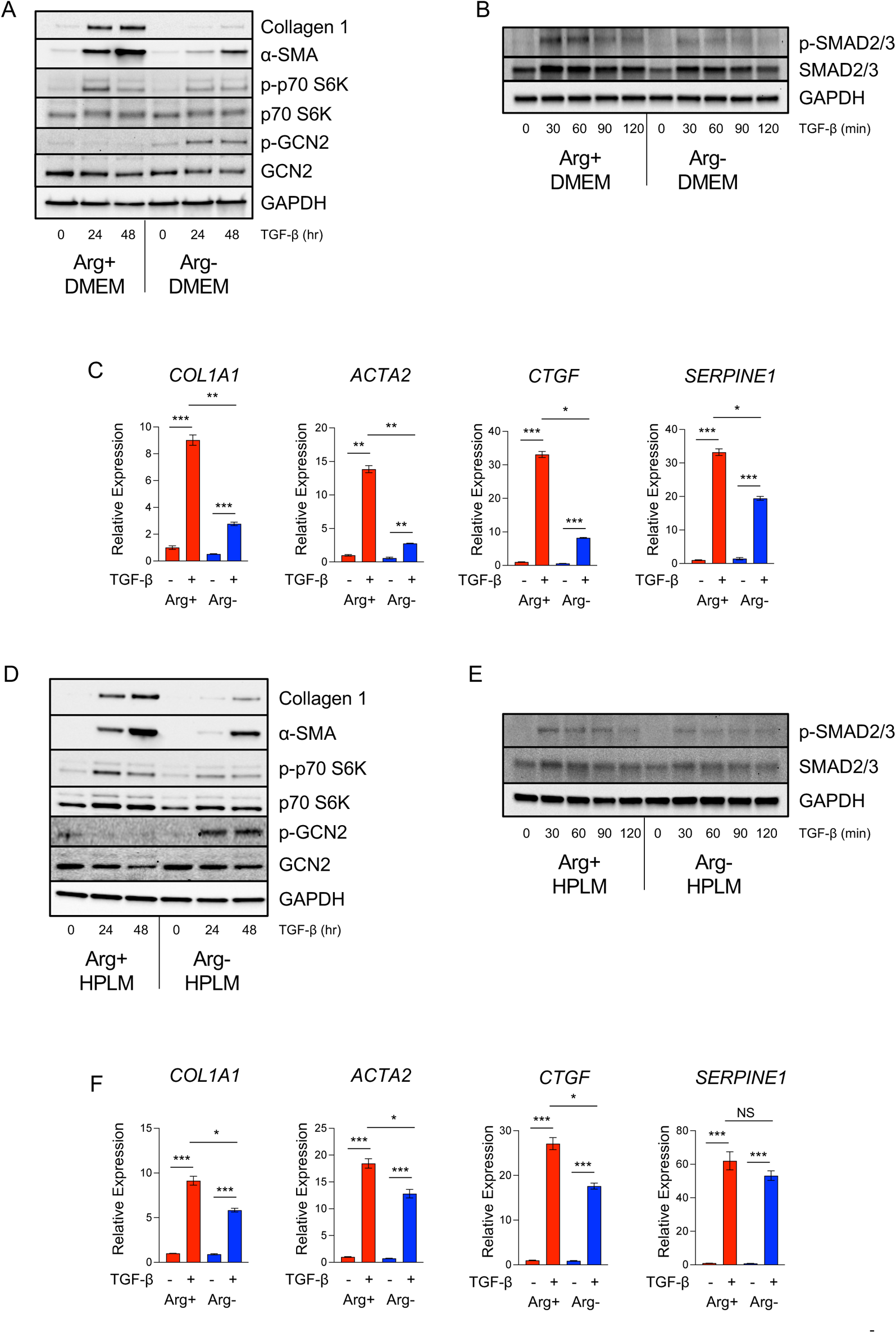
Arginine is required for TGF-β-induced signaling and gene expression in human lung fibroblasts. **(A)** Western blot analysis of collagen 1 and α-smooth muscle actin protein expression and S6-kinase and GCN2 phosphorylation in HLFs cultured in DMEM that contains either 0.4mM arginine or no arginine. Cells were treated with TGF-β for the indicated intervals. **(B)** Western blot analysis of SMAD2/3 phosphorylation in HLFs cultured in DMEM that contains either 0.4mM arginine or no arginine. Cells were treated with TGF-β for the indicated intervals. **(C)** qPCR analysis of *COL1A1*, *ACTA2*, *CTGF*, and *SERPINE1* mRNA expression in HLFs cultured in the DMEM that contains either 0.4mM arginine or no arginine. Cells were treated with TGF-β for 24 hours or left untreated. **(D)** Western blot analysis of collagen 1 and α-smooth muscle actin protein expression and S6-kinase and GCN2 phosphorylation in HLFs cultured in HPLM that contains either 0.11mM arginine or no arginine. Cells were treated with TGF-β for the indicated intervals. **(E)** Western blot analysis of SMAD2/3 phosphorylation in HLFs cultured in HPLM that contains either 0.11mM arginine or no arginine. Cells were treated with TGF-β for the indicated intervals. **(F)** qPCR analysis of *COL1A1*, *ACTA2*, *CTGF*, and *SERPINE1* mRNA expression in HLFs cultured in the HPLM that contains either 0.11mM arginine or no arginine. Cells were treated with TGF-β for 24 hours or left untreated. \**P*<0.05, \*\**P*<0.01, \*\*\**P*<0.001.

Arginine is a conditionally nonessential amino acid, which can be produced *de novo* from its precursors, ornithine and citrulline (**Fig. S1**). Recent development of media formulations that mimic concentrations of metabolites in human plasma has revealed that some cellular phenotypes are dependent on non-physiologic nutrient concentrations of standard culture media such as DMEM [37; 45]. We thus cultured HLFs in Human Plasma Like Medium (HPLM) which contains a reduced concentration of arginine compared to DMEM (0.11mM) but contains the arginine precursors ornithine (0.07mM) and citrulline (0.04mM). HLFs cultured in HPLM had reduced levels of intracellular arginine compared with DMEM-cultured cells but had higher levels of ornithine and citrulline (**Fig. S2A**). We found that HLFs cultured in HPLM exhibited TGF-β-induced production of collagen and α-SMA protein that was similar to that from cells cultured in DMEM (**Fig. S2B**). Sequencing of RNA from DMEM- and HPLM-cultured HLFs showed that TGF-β-induced transcriptional responses were similar between cells cultured in the two media (**Fig. S2C, S2D**). To determine whether cells cultured in HPLM maintained sensitivity to extracellular arginine, we formulated HPLM lacking arginine and analyzed TGF-β-induced protein expression. We found that cells cultured in HPLM maintained sensitivity to extracellular arginine, including GCN2 phosphorylation and loss of collagen induction in the absence of arginine as well as reduced SMAD2/3 phosphorylation (**Fig. 1D, 1E**); however, TGF-β-induced transcription was less sensitive to arginine deprivation in cells cultured in HPLM compared with cells cultured in DMEM. (**Fig. 1F**). These findings suggest that arginine promotes TGF-β-induced signaling and collagen production in HLFs, and that the effect of arginine deficiency can be reduced in the presence of arginine precursors.

### Exogenous citrulline can rescue fibroblast activation in arginine-deprived HLFs

Because cells cultured in HPLM exhibited reduced sensitivity to arginine deprivation, we sought to determine whether this finding was dependent on the presence of ornithine and citrulline in the medium. We thus formulated HPLM lacking, ornithine, and citrulline, and then added back each amino acid individually to assess their role in promoting fibroblast activation downstream of TGF-β. We found that in the absence of all three amino acids (Arg^−^Orn^−^Cit^−^), TGF-β-induced production of collagen and α-SMA was abolished (**Fig. 2A**). Addition of arginine to this media (Arg^+^Orn^−^Cit^−^) resulted in a complete rescue of collagen and α-SMA protein expression. This was associated with increased S6-kinase phosphorylation and loss of GCN2 phosphorylation. Addition of citrulline (Arg^−^Orn^−^Cit^+^), but not ornithine (Arg^−^Orn^+^Cit^−^) partially rescued the effect of arginine deficiency. These findings suggest that while ornithine and citrulline do not play a major role in supporting TGF-β-induced activation of lung fibroblasts when arginine is present, citrulline, but not ornithine can be used as a substrate to promote fibroblast activation in the absence of arginine. Consistent with this, we found that when HLFs were cultured in the presence of ornithine and citrulline, arginine concentrations could be reduced to as low as 0.01mM and TGF-β-mediated induction of collagen and α-SMA protein expression was unaffected (**Fig. 2B**). When cells were cultured in media lacking ornithine and citrulline, arginine concentrations could only be reduced to 0.02mM before expression of collagen and α-SMA protein was inhibited (**Fig. 2C**).

**Figure 2.**
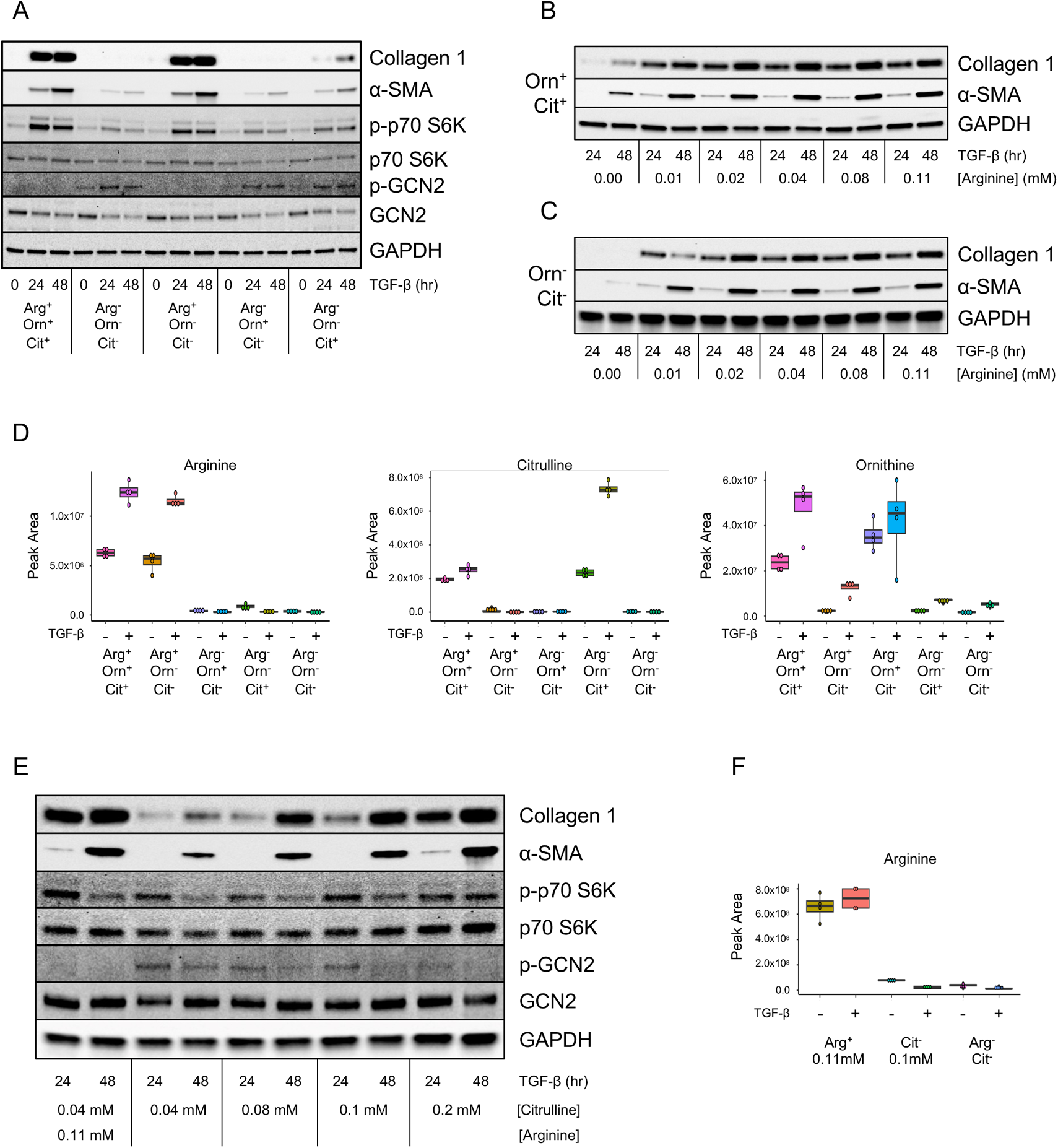
Extracellular citrulline can rescue the effect of arginine depletion in HLFs. **(A)** Western blot analysis of collagen 1 and α-smooth muscle actin protein expression and S6-kinase and GCN2 phosphorylation in HLFs cultured in HPLM that 1) contains arginine, ornithine, and citrulline, 2) contains no arginine, ornithine, or citrulline, 3) contains just arginine, 4) contains just ornithine, 5) contains just citrulline. Cells were treated with TGF-β for the indicated intervals. **(B)** Western blot analysis of collagen 1 and α-smooth muscle actin protein expression and S6-kinase and GCN2 phosphorylation in HLFs cultured in HPLM containing ornithine, citrulline, and the indicated concentrations of arginine. Cells were treated with TGF-β for the indicated intervals. **(C)** Western blot analysis of collagen 1 and α-smooth muscle actin protein expression and S6-kinase and GCN2 phosphorylation in HLFs cultured in HPLM containing no ornithine or citrulline, with the indicated concentrations of arginine. Cells were treated with TGF-β for the indicated intervals. **(D)** Intracellular levels of arginine, ornithine, and citrulline from cells cultured in HPLM as in (A). Cells were treated with TGF-β or left untreated for 48 hours. **(E)** Western blot analysis of collagen 1 and α-smooth muscle actin protein expression and S6-kinase and GCN2 phosphorylation in HLFs cultured in HPLM containing no ornithine or arginine, with the indicated concentrations of citrulline. Cells were treated with TGF-β for the indicated intervals. **(F)** Intracellular levels of arginine from cells cultured in HPLM as in (E). Cells were treated with TGF-β or left untreated for 48 hours.

When we examined intracellular levels of arginine metabolites in HLFs cultured in these media, we found that arginine levels were similar between Arg^+^Orn^+^Cit^+^ cultured cells and Arg^+^Orn^−^Cit^−^ cultured cells, suggesting that minimal arginine synthesis occurs when extracellular arginine is present (**Fig. 2D**). Intracellular ornithine levels were highest in cells cultured in the presence of extracellular ornithine; however, extracellular arginine (Arg^+^Orn^−^Cit^−^) could partially rescue ornithine levels (**Fig. 2D**). Citrulline levels were similar at baseline between Arg^+^Orn^+^Cit^+^ cultured cells and Arg^−^Orn^−^Cit^+^; however intracellular citrulline was highest after TGF-β in cells cultured in only citrulline, suggesting that citrulline uptake is increased in the absence of arginine (**Fig. 2D**). Extracellular citrulline but not ornithine increased arginine levels at baseline (**Fig. 2D**, **S3A**); however, after treatment with TGF-β, intracellular levels of arginine were not significantly different from cells cultured in Arg^−^Orn^−^Cit^−^ media, suggesting that TGF-β increases arginine usage and that 0.04mM extracellular citrulline cannot contribute to arginine accumulation in the absence of extracellular arginine.

We thus sought to determine whether increasing extracellular citrulline could rescue fibroblast activation in the absence of arginine by titrating extracellular citrulline levels in medium lacking arginine and ornithine. We found that increasing extracellular citrulline concentrations to 0.1mM rescued TGF-β-induced collagen and α-SMA protein production (**Fig. 2E**). This was associated with reduced GCN2 phosphorylation on day 2 of TGF-β treatment. Intracellular arginine levels remained very low in cells cultured in 0.1mM extracellular citrulline; however, they were significantly elevated compared to cells cultured in Arg^−^Orn^−^Cit^−^ media (**Fig. 2F**, **S3B**). This suggests while arginine levels were extremely low in the absence of extracellular arginine, *de novo* arginine production was sufficient to maintain the signaling roles of arginine, allowing collagen induction downstream of TGF-β.

### ASS1 promotes fibroblast activation in the absence of arginine

While extracellular citrulline was insufficient to rescue intracellular arginine concentrations in the absence of extracellular arginine, we found that argininosuccinate levels were similar between cells cultured in Arg^+^Orn^−^ Cit^−^ media and Arg^−^Orn^−^Cit^+^ media (**Fig. 3A**). Argininosuccinate is produced from citrulline by argininosuccinate synthase (ASS1) and can then be further converted to arginine by argininosuccinate lyase (ASL) (**Fig. S1**). To determine whether TGF-β-induced lung fibroblast activation in the presence of only citrulline was dependent on ASS1, we knocked down ASS1 expression using siRNA and treated cells with TGF-β in the presence of Arg^−^Orn^−^Cit^−^ media supplemented with either 0.11mM arginine, 0.1mM citrulline, or neither amino acid. Consistent with a role for ASS1 in promoting arginine synthesis in the absence of extracellular arginine, we found that ASS1 was not required for fibroblast activation when extracellular arginine was present (**Fig. 3B**). However, when cells were cultured in 0.1mM citrulline with no arginine, ASS1 became required to support collagen and α-SMA protein production.

**Figure 3.**
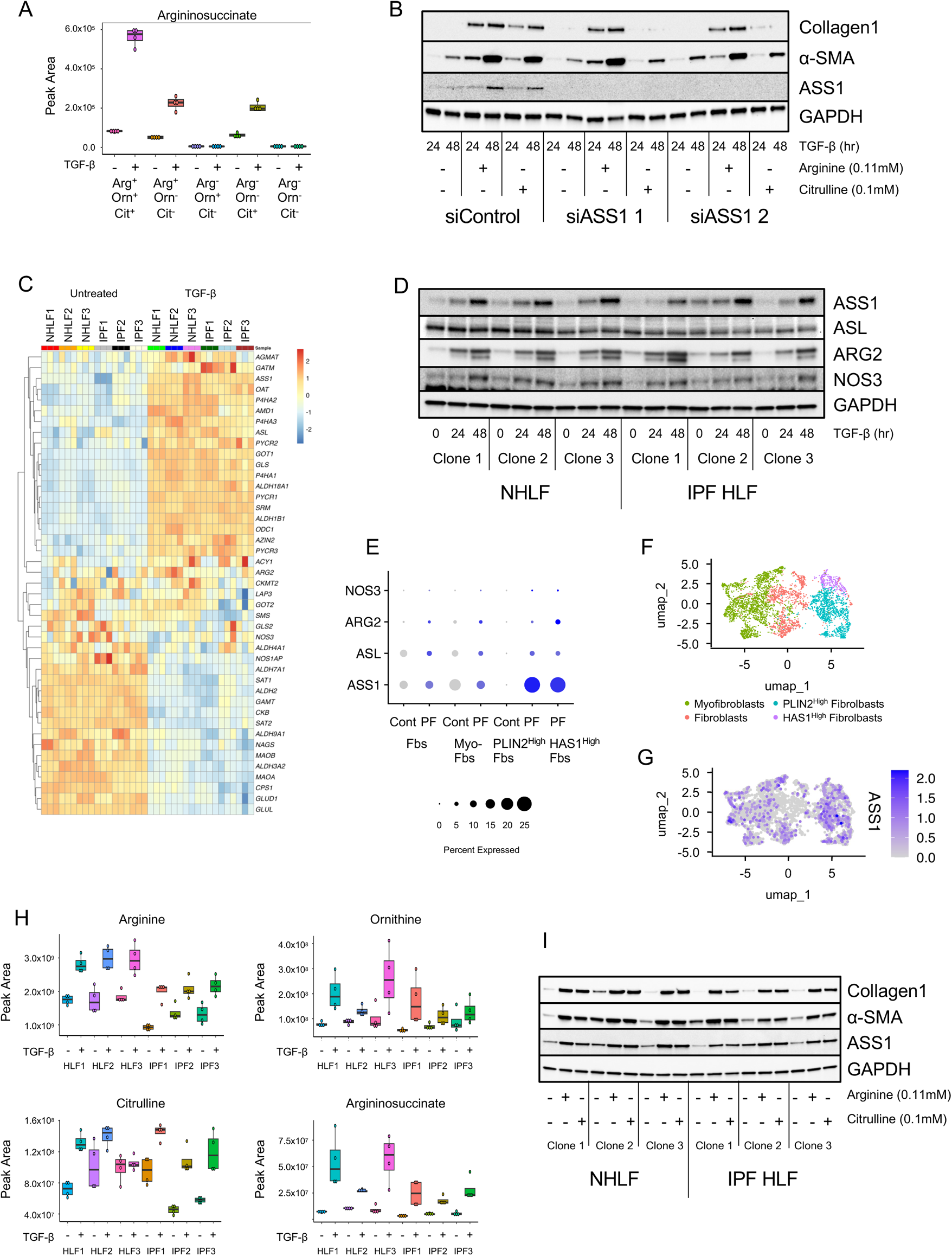
ASS1 promotes de novo arginine production in normal and IPF HLFs. **(A)** Intracellular levels of argininosuccinate from cells cultured in HPLM as in (2A). Cells were treated with TGF-β or left untreated for 48 hours. **(B)** Western blot analysis of collagen 1 and α-smooth muscle actin protein expression in HLFs cultured in HPLM that contains arginine (0.11mM) and citrulline (0.1mM) as indicated. Cells were treated with TGF-β for the indicated intervals. **(C)** Heatmap analysis showing the KEGG arginine and proline metabolism pathway on RNAseq data from 3 clones of normal and 3 clones of IPF HLFs. **(E)** Dot plot representation of the expression of arginine metabolic enzymes in fibroblast subpopulations from pulmonary fibrosis patients and control donor lungs and stratified by disease state as defined by Habermann et al [44]. **(F-G)** UMAP projections of fibroblast subpopulations and ASS1 mRNA expression in fibroblasts from pulmonary fibrosis patients. **(H)** Intracellular levels of arginine, ornithine, citrulline, and argininosuccinate from normal and IPF HLFs cultured in HPLM. Cells were treated with TGF-β or left untreated for 48 hours. **(I)** Western blot analysis of collagen 1 and α-smooth muscle actin protein expression in HLFs cultured in HPLM that contains arginine (0.11mM) and citrulline (0.1mM) as indicated. Cells were treated with TGF-β for the indicated intervals.

ASS1 has been suggested to be reduced in IPF fibroblasts, leading to greater sensitivity of these cells to arginine depletion [22]. Using RNAseq, we queried the KEGG arginine and proline metabolism pathway and compared expression of arginine metabolic enzymes in 3 clones of normal HLFs and 3 clones of IPF HLFs (**Fig. 3C**). We found that TGF-β increased the expression of *ASS1* and *ASL* in both normal and IPF HLFs, and that no significant differences in expression were seen between disease and normal cells. We similarly did not detect any differences in TGF-β-induced protein expression of *ASS1* or *ASL* between disease and normal cells (**Fig. 3D**).

We next analyzed the expression of arginine metabolic enzymes in fibroblast populations from a previously published single cell RNAseq data from pulmonary fibrosis patients and control donors [44]. We found that ASS1 is expressed in a greater percentage of lung fibroblasts than other arginine metabolic enzymes. We also found that expressions of ASS1 and ASL were increased in fibroblasts from patients with pulmonary fibrosis (**Fig. 3E**). ASS1 expression was highest in disease-specific PLIN2^High^ and Has1^High^ fibroblast populations as defined by Habermann *et al*; however, expression was detected in fibroblast and myofibroblast populations as well (**Fig. 3F, 3G**).

Interestingly, even though there were no differences observed between normal HLFs and IPF HLFs in the expression of arginine biosynthetic enzymes, we found that intracellular arginine levels were reduced in IPF HLFs compared with HLFs obtained from non-diseased donors (**Fig. 3H**). Intracellular levels of ornithine and citrulline varied based on clone; however, arginine levels were consistently lower by 30% in IPF HLFs. We thus sought to determine whether IPF HLFs display greater sensitivity to arginine deprivation than normal HLFs. We found that IPF HLFs did not exhibit impaired induction of collagen or α-SMA protein when cultured in Arg^−^Orn^−^ Cit^+^ medium containing 0.1mM citrulline. These findings suggest that despite having reduced intracellular arginine levels compared with control cells, IPF HLFs have no defect in their ability to synthesize arginine.

### Arginine is not significantly catabolized in HLFs

In addition to arginine synthesis, arginine catabolism has also been suggested to play a role in lung fibroblasts in the context of fibrosis; however, metabolic labeling experiments have not been conducted to determine how arginine is used in HLFs. We thus cultured HLFs in HPLM containing ^13^C_6_ Arginine to determine how arginine is catabolized after TGF-β (**Fig. 4A**). Strikingly, despite achieving over 80% labeling of intracellular arginine, we found that neither ornithine nor citrulline were significantly labeled downstream of arginine (**Fig. 4B**, **S4A**). We did detect significant labeling (>40%) in dimethylarginine (**Fig. 4B**, **S4A**). Dimethylarginine is the product of hydrolysis of proteins in which arginine residues have been post-translationally methylated by protein arginine methyltransferases (PRMTs). This finding demonstrates the incorporation of labeled arginine into proteins.

**Figure 4.**
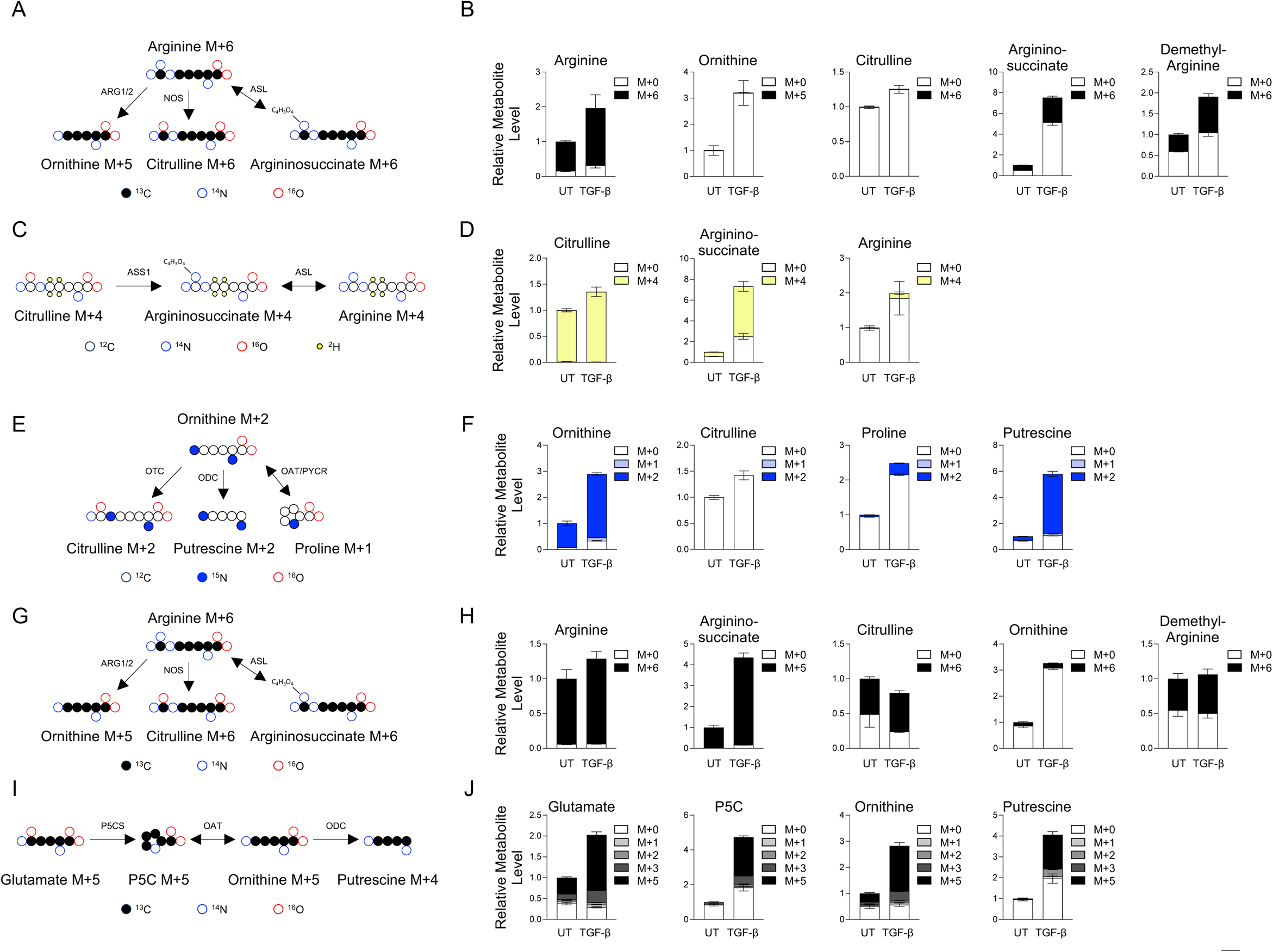
Metabolic tracing of arginine metabolism in human lung fibroblasts. **(A)** Schematic representation of the metabolism of ^13^C_6_ arginine. **(B)** Analysis of cellular arginine, ornithine, citrulline, argininosuccinate, and dimethylarginine in HLFs after labeling with ^13^C_6_ arginine HPLM in the presence or absence of TGF-β. **(C)** Schematic representation of the metabolism of 4,4,5,5-D_4_ citrulline. **(D)** Analysis of cellular citrulline, argininosuccinate, and arginine in HLFs after labeling with 4,4,5,5-D_4_ citrulline HPLM in the presence or absence of TGF-β. **(E)** Schematic representation of the metabolism of ^15^N_2_ ornithine. **(F)** Analysis of cellular ornithine, citrulline, proline, and putrescine in HLFs after labeling with ^15^N_2_ ornithine HPLM in the presence or absence of TGF-β. **(G)** Schematic representation of the metabolism of ^13^C_6_ arginine. **(H)** Analysis of cellular arginine, ornithine, citrulline, argininosuccinate, and dimethylarginine in HLFs after labeling with ^13^C_6_ arginine DMEM in the presence or absence of TGF-β. **(I)** Schematic representation of the metabolism of ^13^C_5_ glutamine. **(J)** Analysis of cellular glutamate, pyrroline-5-carboxylate, ornithine, and putrescine in HLFs after labeling with ^13^C_5_ glutamine DMEM in the presence or absence of TGF-β.

Surprisingly, we also detected significant labeling (>30%) in argininosuccinate, suggesting that ASL is functioning in the reverse direction in lung fibroblasts (**Fig. 4B**, **S4A**). Because argininosuccinate labeled from ^13^C_6_ arginine is labeled on 6 carbons either when produced from reverse ASL reaction or after NOS-dependent citrulline production followed by conversion to argininosuccinate by ASS1, we could not use labeling with ^13^C_6_ arginine to determine which of these pathways was responsible for argininosuccinate production. We thus cultured HLFs in arginine labeled on the two nitrogens of the guanido group (guanido-^15^N_2_). In this case, argininosuccinate produced by the reverse ASL reaction maintains both labeled nitrogens, while one of the nitrogen atoms is lost in the NOS reaction (**Fig. S5A**). Consistent with the reverse ASL reaction, the high degree of labeling on argininosuccinate was M+2 (**Fig. S5B, S5C**).

Because neither ornithine nor citrulline was significantly labeled downstream of arginine, we sought to determine if intracellular levels of these amino acids are determined primarily by their uptake from the extracellular space and whether they are metabolized following uptake. We thus labeled HLFs in HPLM containing either ornithine labeled on both nitrogen atoms (^15^N_2_), or citrulline labeled with deuterium atoms on the 4 and 5 carbons (4,4,5,5-D_4_) (**Fig. 4C, 4E**). We found that intracellular citrulline in HLFs comes almost 100% from extracellular citrulline (**Fig. 4D**, **S4B**). Labeling from citrulline was found on argininosuccinate, demonstrating that most argininosuccinate in HLFs comes from exogenous citrulline. A small portion (<10%) of arginine was also labeled downstream of citrulline (**Fig. 4D**, **S4B**). This finding suggests that the ASL reaction may function in either direction in HLFs depending on cellular conditions or metabolite concentrations.

Labeling HLFs with ornithine also showed that most intracellular ornithine comes from extracellular ornithine (**Fig. 4F**, **S4C**). We did not detect any labeling from ornithine on citrulline, demonstrating that HLFs lack a complete urea cycle. A small amount of labeling was found on proline, while the majority of the polyamine putrescine was labeled by extracellular ornithine (**Fig. 4F**, **S4C**). These results suggest that a major role of extracellular ornithine in lung fibroblasts is polyamine production.

Because we were unable to detect arginine catabolism in HLFs cultured in HPLM, we considered the possibility that extracellular ornithine and citrulline inhibit the catabolism of arginine under these conditions. We thus cultured cells in DMEM containing ^13^C_6_ arginine to determine how arginine is metabolized in the absence of its extracellular precursors (**Fig. 4G**). We found that culturing cells in DMEM resulted in the majority of citrulline being labeled on 6 carbons, indicating that NOS is active when HLFs are cultured in DMEM (**Fig. 4H**, **S4D**). Surprisingly, only a small amount (<15%) of ornithine and downstream putrescine was labeled from arginine, indicating a low amount of arginase activity (**Fig. 4H, S6A, S6B**). No proline was found to be labeled from arginine (**Fig. S6A, S6B**). We also found that argininosuccinate was almost completely labeled M+6 by ^13^C_6_ arginine (**Fig. 4H**, **S4D**). Labeling with guanido-^15^N_2_ arginine demonstrated that this occurred through the reverse ASL reaction (**Fig. S5D, S5E**).

Arginine is predicted to be the major source of cellular ornithine and polyamines in DMEM-cultured cells; thus, it was surprising that the majority of these molecules were unlabeled after culture in ^13^C_6_ arginine. The OAT reaction which produces pyrroline-5-carboxylate (P5C), a precursor for proline, is reversible. We and others have shown that glutamate-dependent production of P5C and proline through ALDH18A1 is increased in lung fibroblasts downstream of TGF-β [15; 46]. We thus sought to determine whether glutamate is a major source of ornithine and polyamines through the reverse OAT reaction. We thus labeled HLFs with ^13^C_5_ glutamine (**Fig. 4I**). We found that this resulted in the majority of glutamate being labeled through the glutaminase reaction (**Fig. 4J**, **S4E**). As we have previously demonstrated, glutamine is an important source of P5C and proline in HLFs (**Fig. 4J, S6C, S6D**). We also found significant labeling from glutamine on ornithine and putrescine, demonstrating that glutamine, and not arginine is the major source of these metabolites in HLFs cultured in DMEM (**Fig. 4J**, **S4E**).

## DISCUSSION

Metabolic reprogramming in lung fibroblasts is required to support matrix protein production in the context of pulmonary fibrosis [9–12]. Arginine and its metabolism have been linked with matrix production in lung fibroblasts; however, a thorough analysis of the role of arginine biosynthesis and catabolism has not previously been conducted. Here, our work suggests that the major role of arginine in lung fibroblasts is to support production of arginine-containing proteins and to signal amino acid sufficiency to mTOR and GCN2. Surprisingly, catabolism of arginine does not occur to a significant extent in HLFs.

When HLFs were cultured in medium formulated to mimic human plasma, we found that labeling from extracellular arginine only significantly accumulated on dimethylarginine and on argininosuccinate. Dimethylarginine is a product of the hydrolysis of proteins containing arginine residues that were post-translationally methylated; thus, we provide evidence of the use of extracellular arginine the charging of arginyl-tRNAs. Our results suggest that the ASL reaction in HLFs proceeds primarily in the direction of argininosuccinate synthesis from arginine. ASL cleaves argininosuccinate, producing arginine and fumarate. The ASL reaction is known to be reversible and is a net producer of argininosuccinate in certain conditions such as in fumarate hydratase-deficient cancers, in which the consumption of fumarate by ASL enables survival [47]. Whether the reverse ASL reaction plays an important role in lung fibroblasts remains to be determined.

While we did not observe significant arginine catabolism in cells cultured in HPLM, we found evidence that in the absence of extracellular ornithine and citrulline, both arginase and NOS-dependent pathways are active in HLFs, leading to M+6 labeling on citrulline (NOS) and M+5 labeling on ornithine (arginase). While it is not possible to determine from our data if either pathway is preferentially used by HLFs, it is striking that the minority of cellular ornithine comes from arginase activity when HLFs are cultured in DMEM. We and others have previously demonstrated that glutamine-dependent proline synthesis is required to support collagen production downstream of TGF-β [15; 46]. Ornithine has been hypothesized to be a precursor for proline through the activity of OAT; however, our findings suggest that OAT activity proceeds primarily in the direction of ornithine synthesis even in cells cultured in DMEM which contains abundant arginine and no proline. OAT expression has been shown to be elevated in IPF lung tissue and expressed in fibroblasts. Inhibition of OAT reduces collagen protein production downstream of TGF-β [23; 29]. Our results suggest that this may be more related to the production of ornithine and downstream production of polyamines than to production of proline to support collagen synthesis. It is also possible that a nonmetabolic role for OAT exists.

Our results show that in the absence of extracellular citrulline, most intracellular citrulline comes from arginine metabolism through NOS. Little is known about the role of NOS in lung fibroblasts or in IPF. Alveolar NO concentrations are elevated in IPF patients [48]; however it is not known what cell types are the main contributor to this elevation. NOS2 expression is increased when lung fibroblasts are induced to proliferate and NOS inhibition reduced fibroblast growth rate [49]. Some evidence suggests that NO production in fibroblasts promotes collagen synthesis [50–52]; however, NOS triple knockout mice (lacking all 3 NOS isoforms) exhibit increased severity of pulmonary fibrosis after instillation of bleomycin [53]. Our transcriptomic findings suggest that NOS3 is the primary NOS expressed in lung fibroblasts both *in vitro* and *in vivo*; however, our metabolic labeling experiments suggest that it does not play a major role in arginine metabolism.

ASS1 has been suggested to be reduced in IPF HLFs, increasing their sensitivity to arginine depletion. Our findings show that ASS1 is required to support collagen production when HLFs are cultured in the absence of arginine. We found no evidence that ASS1 was limiting in IPF HLFs as these cells could produce collagen when cultured using citrulline as a precursor for arginine production. We did note that arginine concentrations were reduced in IPF HLFs. Whether this is due to increased arginine catabolism, reduced arginine uptake, or reduced breakdown of arginine-containing proteins will need to be explored further. While HPLM better mimics the concentrations of amino acids found in human plasma, it is unknown what the local nutrient concentrations are in fibrotic lungs. While our findings demonstrate that reducing arginine levels to one-fifth of what is found in human plasma does not affect the ability of HLFs to support collagen production, even in the absence of extracellular citrulline, production of arginine may be required if local arginine levels are depleted. Similarly, the local concentrations of ornithine and glutamine may affect the contribution of these amino acids to polyamine production.

Our work shows that TGF-β-induced collagen production can occur at very low levels of intracellular arginine and that intracellular levels of arginine above a threshold required to activate mTORC1 and prevent GCN2 activation is sufficient to support collagen protein production. Surprisingly, we found that arginine depletion led to reduced SMAD phosphorylation and SMAD-dependent transcription downstream of TGF-β. We have previously shown that SMAD phosphorylation occurs independently of PI3K and mTOR signaling [30]. Our findings suggest that GCN2 may feed back onto the TGF-β pathway, potentially acting as both an inhibitor of protein translation and as an inhibitor of profibrotic signaling pathways under amino acid-deficient conditions.

In conclusion, we find that arginine is a major regulator of TGF-β-induced activation of lung fibroblasts. While arginine biosynthesis and catabolism can be increased depending on local nutrient conditions, our findings suggest that the major requirements for arginine are for production of arginine-containing proteins and for its role as a signaling molecule. Our findings suggest that therapies that target the signaling role of arginine may prevent collagen production by activated tissue fibroblasts in IPF and in other fibrotic diseases.

## Author contributions

**Robert Hamanaka:** Conceptualization, Data Curation, Formal Analysis, Funding Acquisition, Investigation, Methodology, Supervision, Writing-Original Draft, Writing-Review and Editing. **Kun Woo Shin:** Data Curation, Formal Analysis, Investigation, Writing-Review and Editing. **Volkan Atalay:** Formal Analysis, Investigation, Writing-Review and Editing. **Rengul Cetin-Atalay:** Formal Analysis, Investigation, Writing-Review and Editing. **Hardik Shah:** Data Curation, Formal Analysis, Investigation, Methodology, Writing-Review and Editing. **Jennifer Szafran:** Conceptualization, Investigation. **Parker Woods:** Investigation. Angelo Meliton: Investigation. **Obada Shamaa:** Investigation. **Yufeng Tian:** Investigation, Data Curation. **Takugo Cho:** Investigation, Data Curation. **Gökhan M. Mutlu:** Conceptualization, Formal Analysis, Funding Acquisition, Methodology, Supervision, Writing-Review and Editing.

---

**Figure S1.**
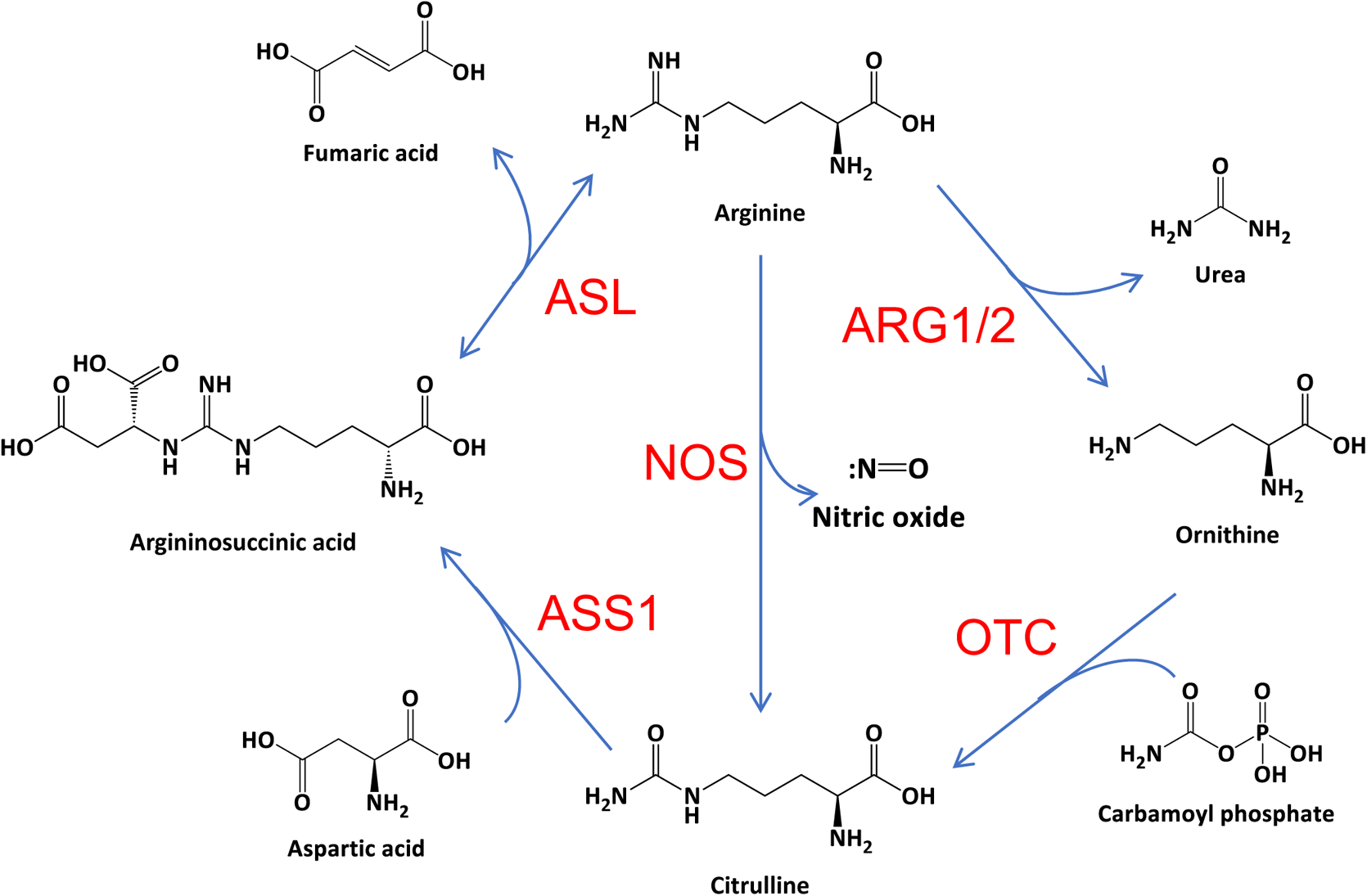
Metabolism of arginine. Schematic illustration of arginine metabolism. Arginine is catabolized by arginase (ARG1, ARG2) or by nitric oxide synthase (NOS), producing urea or nitric oxide, respectively. In cells with a functional urea cycle, ornithine is converted to citrulline by ornithine transcarbamylase (OTC). Citrulline is can be converted back to arginine through the combined actions of argininosuccinate synthase 1 (ASS1) and argininosuccinate lyase (ASL).

**Figure S2.**
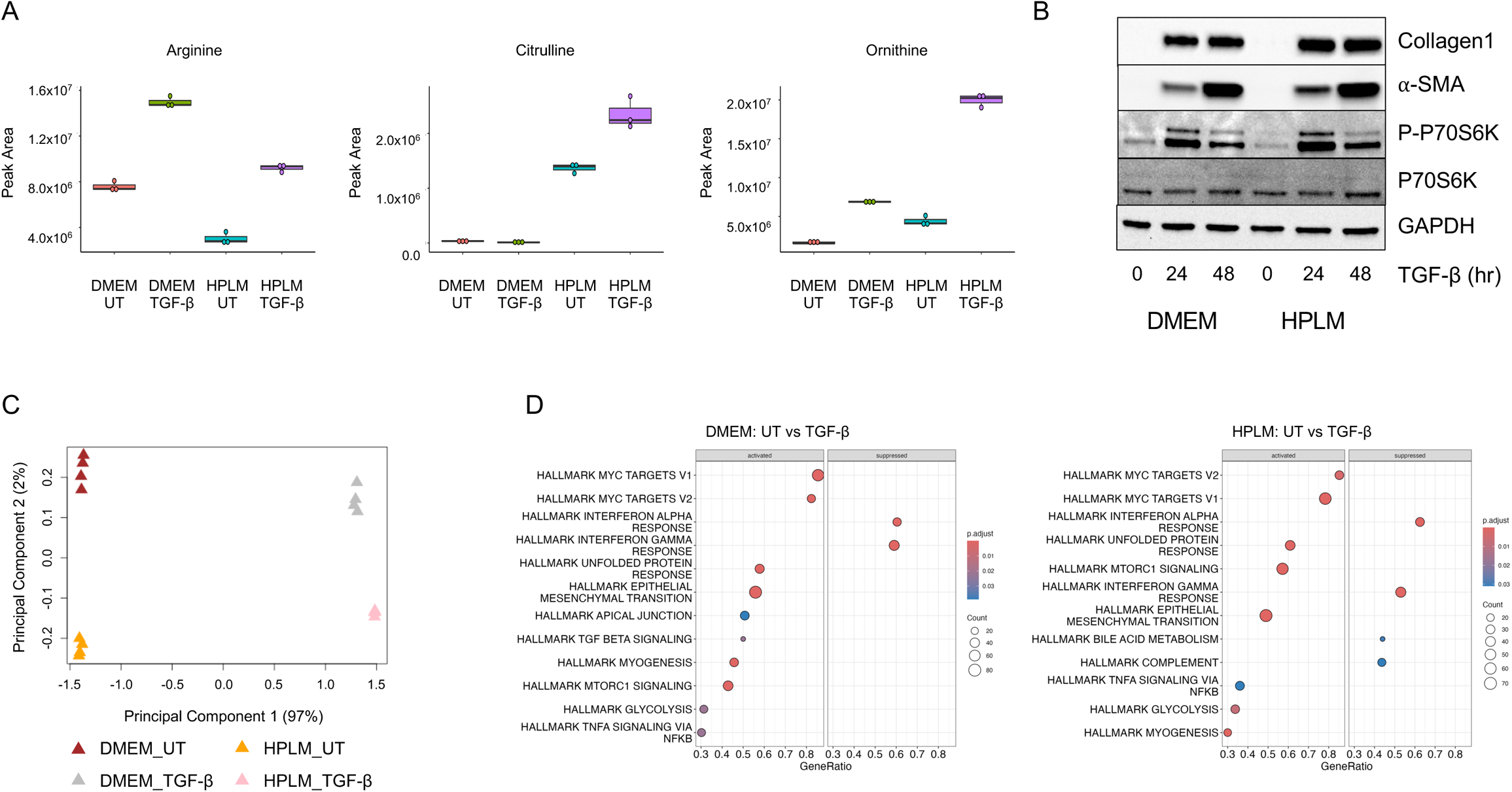
Comparison of cellular metabolites and gene expression in HLFs cultured in DMEM or HPLM. **(A)** Intracellular levels of arginine, citrulline, and ornithine in HLFs cultured in DMEM or HPLM and treated with TGF-β or left untreated for 48 hours. **(B)** Western blot analysis of collagen 1 and α-smooth muscle actin protein expression and S6-kinase phosphorylation in HLFs cultured in either DMEM or HPLM. Cells were treated with TGF-β for the indicated intervals. **(C)** Multidimensional scaling plot of differentially expressed genes (DEGs) in HLFs cultured either DMEM or HPLM and treated with TGF-β or left untreated. **(D)** Significantly activated and suppressed Molecular Signatures Database (MSigDB) Hallmark pathways enriched in DEGs between untreated and TGF-β-treated HLFs. Cells were cultured in DMEM or HPLM as indicated.

**Figure S3.**
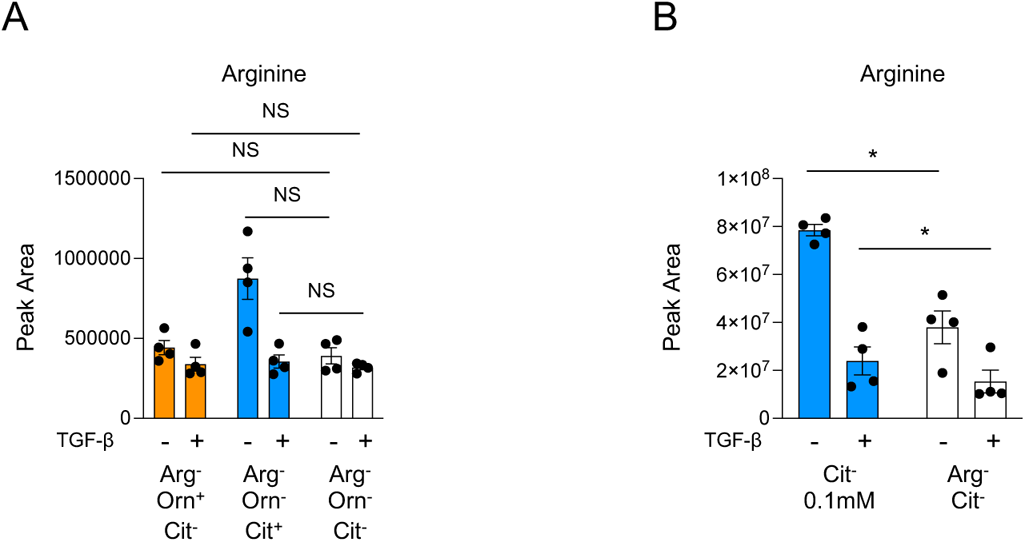
Extracellular citrulline contributes to intracellular arginine levels in the absence of extracellular arginine. **(A)** Intracellular arginine levels in in HLFs cultured in arginine-free HPLM containing either ornithine (0.07mM), citrulline (0.04mM), or neither amino acid. Cells were treated with TGF-β or left untreated for 48 hours. **(B)** Intracellular arginine levels in HLFs cultured in arginine-free HPLM containing either 0.1mM citrulline or 0mM citrulline. \**P*<0.05.

**Figure S4.**
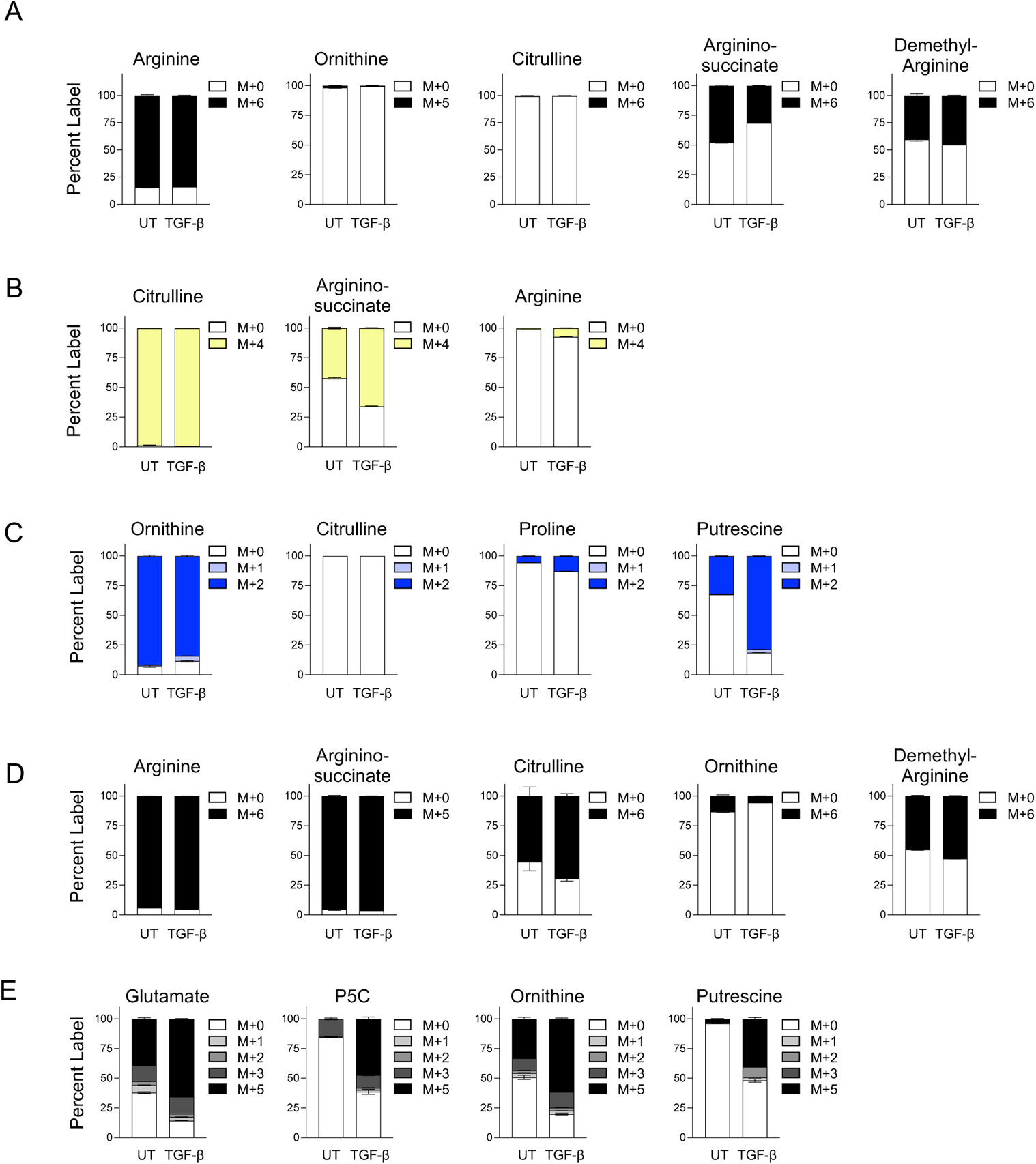
Percent labeling of arginine metabolism in human lung fibroblasts. **(A)** Analysis of cellular arginine, ornithine, citrulline, argininosuccinate, and dimethylarginine in HLFs after labeling with ^13^C_6_ arginine HPLM in the presence or absence of TGF-β. **(B)** Analysis of cellular citrulline, argininosuccinate, and arginine in HLFs after labeling with 4,4,5,5-D_4_ citrulline HPLM in the presence or absence of TGF-β. **(C)** Analysis of cellular ornithine, citrulline, proline, and putrescine in HLFs after labeling with ^15^N_2_ ornithine HPLM in the presence or absence of TGF-β. **(D)** Analysis of cellular arginine, ornithine, citrulline, argininosuccinate, and dimethylarginine in HLFs after labeling with ^13^C_6_ arginine DMEM in the presence or absence of TGF-β. **(E)** Analysis of cellular glutamate, pyrroline-5-carboxylate, ornithine, and putrescine in HLFs after labeling with ^13^C_5_ glutamine DMEM in the presence or absence of TGF-β.

**Figure S5.**
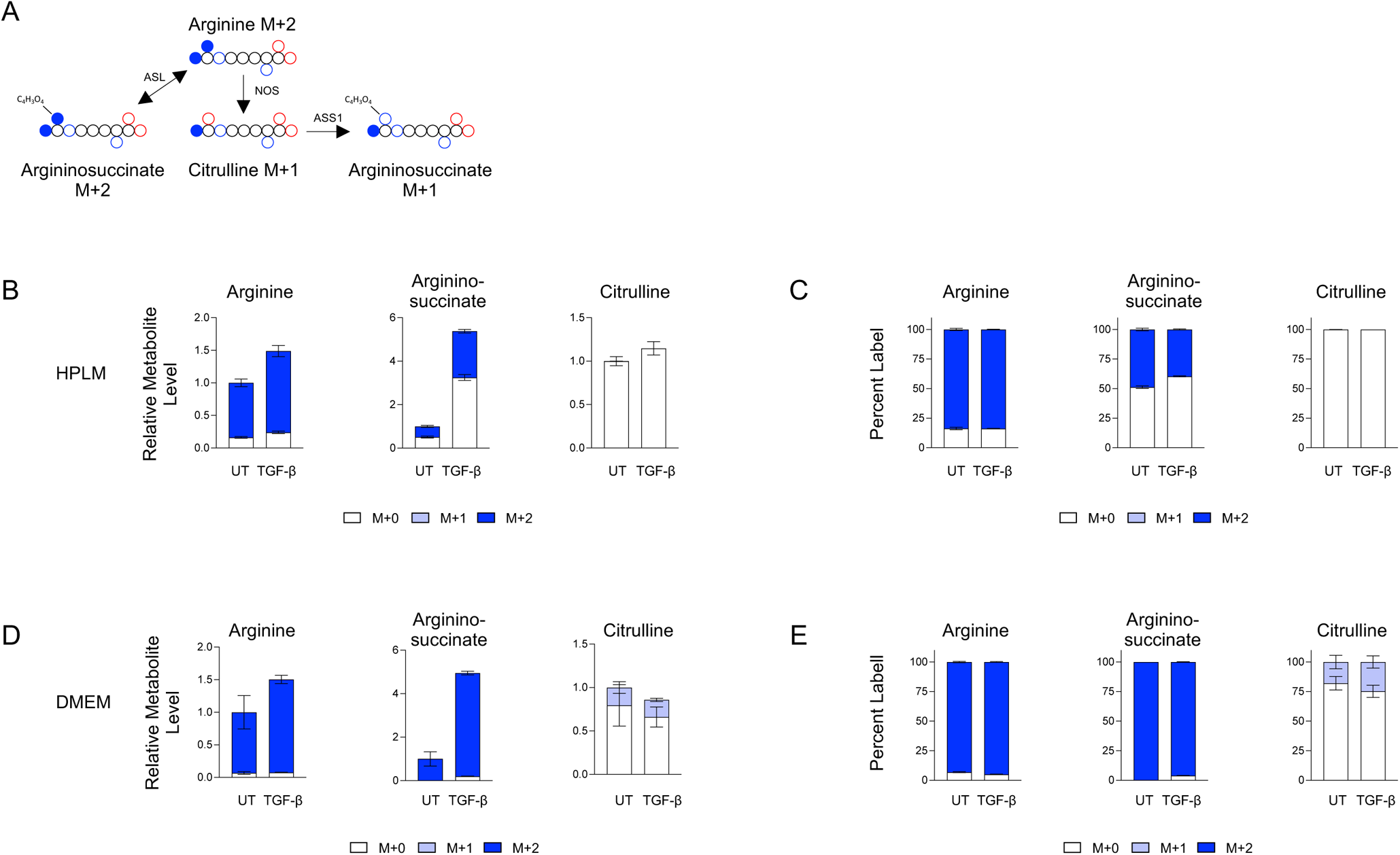
Tracing of guanido-^15^N_2_ arginine in human lung fibroblasts. **(A)** Schematic representation of the metabolism of guanido-^15^N_2_ arginine. **(B)** Analysis of cellular arginine, argininosuccinate, and citrulline in HLFs after labeling with guanido-^15^N_2_ arginine HPLM in the presence or absence of TGF-β. **(C)** Analysis of (B) presented as percent labeling. **(D)** Analysis of cellular arginine, argininosuccinate, and citrulline in HLFs after labeling with guanido-^15^N_2_ arginine DMEM in the presence or absence of TGF-β. **(E)** Analysis of (D) presented as percent labeling.

**Figure S6.**
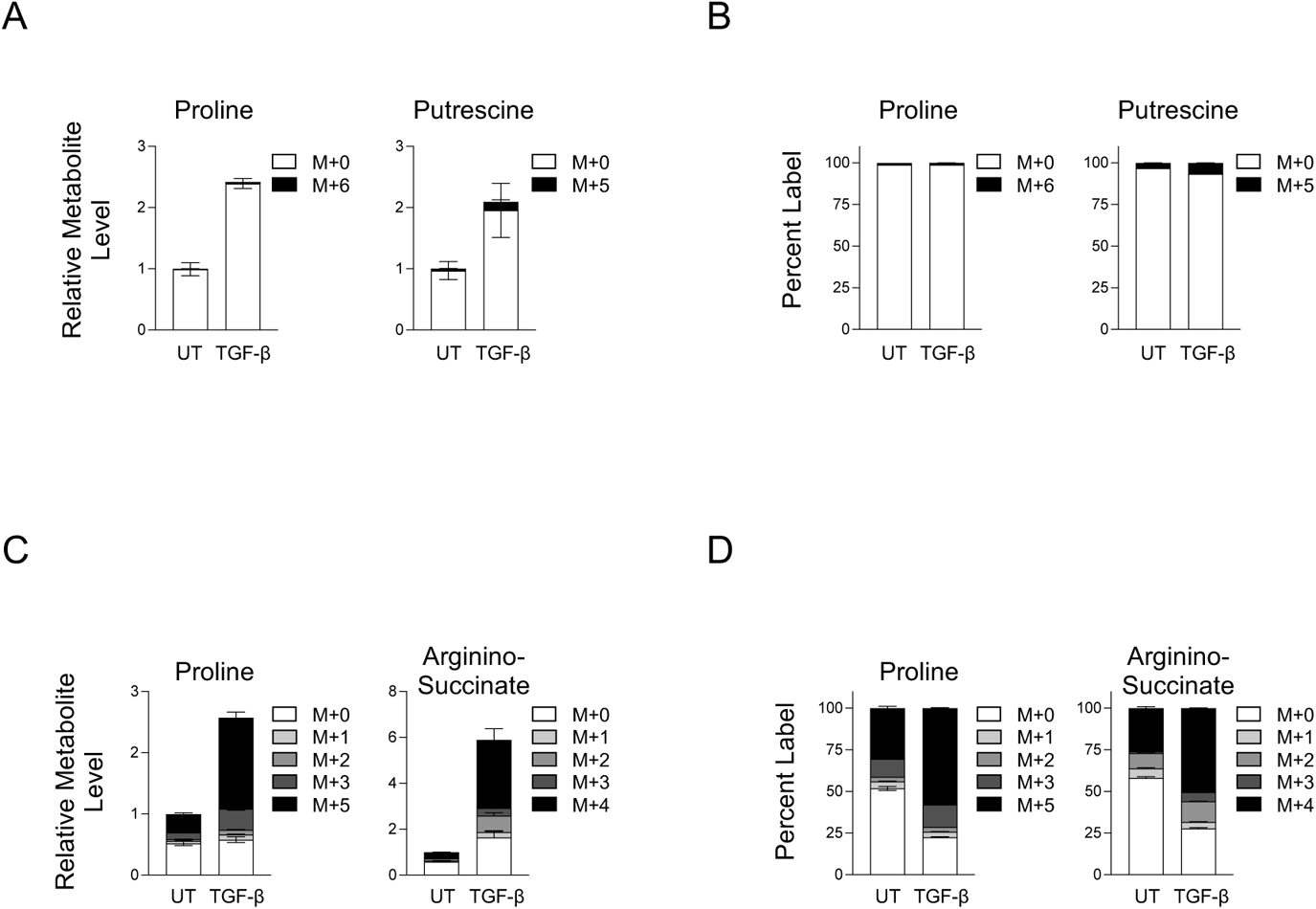
Metabolic tracing of arginine and glutamine in human lung fibroblasts cultured in DMEM. **(A)** Analysis of cellular proline and putrescine in HLFs after labeling with ^13^C_6_ arginine DMEM in the presence or absence of TGF-β. **(B)** Analysis of (A) presented as percent labeling. **(C)** Analysis of cellular proline and argininosuccinate in HLFs after labeling with ^13^C_5_ glutamine DMEM in the presence or absence of TGF-β. **(D)** Analysis of (C) presented as percent labeling.

